# Genomic architecture of artificially and sexually selected traits in a wild cervid

**DOI:** 10.1101/841528

**Authors:** S. J. Anderson, S. D. Côté, J. H. Richard, A. B. A. Shafer

## Abstract

Characterization of the genomic architecture of fitness-related traits such as body size and male ornamentation in mammals provides tools for conservation and management: as both indicators of quality and health, these traits are often subject to sexual and artificial selective pressures. Here we performed high-depth whole genome re-sequencing on pools of individuals representing the phenotypic extremes in our study system for antler and body size in white-tailed deer (*Odocoileus virginianus*). Samples were selected from a tissue repository containing phenotypic data for 4,466 male white-tailed deer from Anticosti Island, Quebec, with four pools representing the extreme phenotypes for antler and body size in the population, after controlling for age. Our results revealed a largely panmictic population, but detected highly diverged windows between pools for both traits with high shifts in allele frequency (mean allele frequency difference of 14% for and 13% for antler and body SNPs in outlier windows). These regions often contained putative genes of small-to-moderate effect consistent with a polygenic model of quantitative traits. Genes in outlier antler windows had known direct or indirect effects on growth and pathogen defence, while body genes, overall GO terms, and transposable element analyses were more varied and nuanced. Through qPCR analysis we validated both a body and antler gene. Overall, this study revealed the polygenic nature of both antler morphology and body size in free-ranging white-tailed deer and identified target loci for additional analyses.

## Introduction

Quantifying the genomic architecture underlying phenotypes in natural populations provides insights into the evolution of quantitative traits (Stinchcombe & Hoekstra, 2008). Some quantitative traits are correlated with metrics of fitness, and as such are particularly important because they might directly influence population viability (Kardos & Shafer, 2018). This relationship between genomic architecture and quantitative traits, however, is not easy to empirically identify (Barghi et al., 2020), and often has unclear and unpredictable responses to selection (Bünger et al., 2005). Outside of a few traits in well studied systems such as horn morphology in Soay sheep (Johnston et al., 2011, 2013) and migration timing in Pacific salmon (Prince et al., 2017), there are few studies that have successfully identified quantitative trait loci (QTL) in wild vertebrate populations with effects on fitness-related traits.

Genome-wide association scans (GWAS) have the potential to identify links between genomic regions and fitness relevant traits that can directly inform management and have become more prominent in non-model organisms (Ellegren 2014; Santure & Grant 2018). The variations of expressed phenotypes that are observed in wild populations are influenced by demographic, environmental and epigenetic factors (Ellegren & Sheldon 2008), which leads to difficulties when trying to quantify the direct influence of genetic polymorphisms, and more generally differentiate the effects of drift from selection (Kardos & Shafer 2018). Most traits also appear to be composed of many genes of small effect, and heritability for complex traits are often modulated by genes outside of core pathways and coding regions (Robinson et al., 2013; Santure et al., 2013; Boyle et al., 2017; Barghi et al. 2020). Population genetic modeling allows disentangling phenotypic plasticity and responses to environmental change, while expanding our understanding of the evolutionary processes in wild populations (Santure & Grant 2018). Where traits directly linked to reproductive success are also those humans target via harvesting (Coltman et al., 2003; Kuparinen & Festa-Bianchet, 2017), there is a need to understand the genomic architecture to predict evolutionary consequences (Kardos & Luikart, 2015), considering that the two modes of selection (natural and artificial) might not always act in concert, and can vary spatially and temporally (Edeline et al. 2007).

For traits relevant to fitness, including those under sexual selection, we might expect directional selection towards the dominant or desired phenotype. In long term studies of wild populations, however, there is often a relatively high degree of observed genetic variation underlying sexually selected traits (Johnston et al., 2013; Berenos et al., 2015; Malenfant et al., 2018), with the proposed mechanism for this genetic variation being balancing selection (Mank, 2017; but see Kruuk et al. 2007). For example, Johnston et al. (2013) showed that variation is maintained at a single large effect locus through a trade-off between observed higher reproductive success for large horns and increased survival for smaller horns in Soay sheep. These trade-offs, creating signatures consistent with balancing selection, are also observed in cases where social dominance structures exist as a result of male-male competition for female mate acquisition, where alternative mating strategies have been suggested as a means for younger or subordinate males to achieve greater mating success (Foley et al., 2015).

### Emerging approaches to investigate the genomic architecture of complex phenotypes

The methodological approach of analyzing extreme phenotypes seeks to sample individuals representing the extreme ends of the spectrum for any observable phenotype, instead of randomly sampling individuals from the entire distribution (Perez-Gracia et al., 2002; Li et al., 2011; Emond et al., 2012; Barnett et al., 2013; Gurwitz & McLeod, 2013; Kardos et al., 2016). Pooled sequencing (pool-seq) uses the resequencing of pooled DNA from individuals within a population and is a cost-effective alternative to the whole genome sequencing of every individual (Anand et al., 2016) with lower coverage options emerging (Tilk et al. 2019). Individual identity is lost through the pooling of DNA, but the resulting allelic frequencies are representative of the population and can be used to conduct standard population genomic analyses, including outlier detection (Schlötterer et al., 2014). Pool-seq methods have been applied with the purposes of identifying genes of moderate-to-large effect loci in a variety of taxa including colour morphs in butterflies and birds, abdominal pigmentation in *Drosophila*, and horn size of wild Rocky Mountain bighorn sheep (Kardos et al, 2015; Endler et al., 2017; Neethiraj et al., 2017).

Here, we explored the genic basis for phenotypic variation by sampling the extreme phenotypes in a non-model big game species, the white-tailed deer (*Odocoileus virginianus*, WTD). Two traits are of particular interest in WTD; body size and antler size, which have a degree of observable variation (Hewitt 2011) and are connected to individual reproductive success (DeYoung et al., 2009; Newbolt et al., 2016; Jones et al., 2018). Heritability for antler and body measurements are moderate to high for antler features and body size (Michel et al., 2016; Williams et al., 1994; Jamieson et al., 2020). Large antlers and body size are also sought after by hunters, both as trophies and food sources throughout North America. Efforts to compare harvested to non-harvested (i.e. naturally shed) antlers suggest the potential for artificial selection with those harvested by hunters being larger in size (Schoenebeck & Petersen 2014 but see Ditchkoff et al. 2000), which is similar to what has been documented in other cervids (Ramanzin & Sturaro, 2014; Pozo et al., 2016; Balčiauskas et al., 2017; García-Ferrer et al., 2019). Our objective was to identify genomic windows associated with variation in antler and body size phenotypes in WTD, under the hypothesis that many genes of small to moderate effect underly these phenotypes, but core pathways should be related to these phenotypes. We performed whole genome re-sequencing on pools representing the extreme distribution of antler and body size phenotypes in a wild, but intensively monitored WTD population.

## Materials and Methods

### Study area

Anticosti Island (49°N, 62°W; 7,943 km^2^) is located in the Gulf of St. Lawrence, Québec (Canada) at the northeastern limit of the white-tailed deer range (Figure 1). The island is within the balsam fir-white birch bioclimatic region with a maritime sub-boreal climate characterized by cool and rainy summers (630mm/year), and long and snowy winters (406 cm/year; Environment Canada, 2006; Simard et al., 2010). The deer population was introduced in 1896 with ca. 220 animals and rapidly increased. Today, densities are >20 deer/km^2^ and can exceed 50 deer/km^2^ locally (Potvin & Breton 2005) with an estimated population >160,000 individuals.

### Sample Collection and Modeling

We collected tissue samples stored in 95% ethanol and phenotypic data on 4,466 male deer harvested by hunters from September to early December, 2002–2014 on Anticosti Island. The sample and measurements were completed as part of a large effort led by the NSERC Industrial Research Chair in integrated resource management of Anticosti Island focussed on collecting data on the body condition of deer on the island. We used cementum layers in incisor teeth to age individuals (Hamlin et al., 2000). Two metrics of antler size were selected: the number of antler tines or points (>2.5 cm) and beam diameter (measured at the base; ±0.02 cm) (Simard et al., 2014). We used one metric for body size: body length which is correlated to other metrics such as body weight and hind foot length (Bundy et al., 1991). Because antler and body size of male cervids are correlated to age (Solberg et al., 2004; Nilsen & Solberg, 2006), we controlled for age before ranking males according to antler and body size. We used linear models to assess the relationship between age and each metric separately. We computed an antler and body size index based on the average rank of each individual’s residual variation for the phenotypic metrics. The top and bottom 150 individuals from each group were selected from the available database for DNA extraction, with only deer having equal representation for large and small phenotypes from the same given year were included in the final pool in an effort to limit temporal variation. This created four groups: large antler (LA), small antler (SA), large body size (LB), and small body size (SB) (Table S1).

### DNA Extraction and Genome Sequencing

We isolated DNA from tissue using the Qiagen DNeasy Blood & Tissue Kit. The concentration of each DNA extract was determined using a Qubit dsDNA HS Assay Kit (Life Technologies, Carlsbad, CA, USA). We aimed to sequence each pool to 50X as per recommendations of Schlötterer et al. (2014). Equal quantities of DNA (100 ng/sample) were combined into representative pools for LA (n=48), and SA (n=48), LB (n=54), and SB (n=61) for a desired final concentration of 20 ng/ul combined DNA for each pool. Sequencing was conducted at The Centre for Applied Genomics (Toronto, ON, Canada) on an Illumina HiSeqX with 150 bp pair-end reads (Table S2); meaning each individual on average should have at least one chromosome sampled.

### Genome Annotation

A newly generated draft WTD genome constructed from long (PacBio) and short-read (Illumina) data was used as a reference (NCBI PRJNA420098; Accession No. JAAVWD000000000). We performed a full genome annotation by masking repetitive elements throughout the genome using a custom WTD database developed through repeat modeler v1.0.11 (Smit & Hubley, 2015) in conjunction with repeatmasker v4.0.7 (Smit, Hubley & Green, 2015) using the NCBI database for artiodactyla, without masking low complexity regions. The masked genome was then annotated using the MAKER2 v2.31.9 pipeline (Holt & Yandell, 2011). We used a three-stage process (Laine et al., 2016) utilizing publicly available white-tailed deer EST and Protein sequences available through NCBI for initial training with SNAP v2013-11-29 (Korf, 2004). A resulting hmm file was generated from the GFF output and was used as evidence for the MAKER2 prediction software, again using SNAP. Evidence from the prior SNAP trails in gff format were used as evidence for the generation of a training data set for the AUGUSTUS v3.3.2 gene prediction software. The final annotation was completed using MAKER2 and AUGUSTUS. Gene IDs were generated using blastp v2.9.0 (Johnson et al., 2008) on the WTD annotation protein transcripts, restricting the blast search to human protein annotations in the uniport database (parameters -max_hsps 1 - max_target_seqs 1 -outfmt 6 -taxids 9606).

### Mapping and characterization of SNPs

We performed initial quality filtering for all reads using fastqc and trimmed reads for quality and adaptors using the default Trimmomatic v.0.36 settings (Bolger et al. 2014). Reads were then aligned to the unmasked WTD reference genome with BWA-mem v0.7.17 (Li, 2013). We used samtools v1.10 (Li et al., 2009) to merge and sort all aligned reads into four files for each representative pool. We then filtered for duplicates using Picard v2.20.6 (Broad Institute, 2020), obtained uniquely mapped reads with samtools, and conducted local realignment using GATKv3.8 (Van der Auwera et al. 2013) prior to calling SNPs. Using samtools we called SNPs with mpileup (parameters = -B -q20). We filtered out the masked regions and indels with 5bp flanking regions and removed all scaffolds <= 50 kb.

Genome wide differences were calculated between large and small phenotypes for antler and body size. We used a sliding window approach with window sizes of 1000 bp with a step size of 500 bp and minimum covered fraction of 0.8 to test for allele frequency differences using Fisher’s exact test (FET) and fixation index (*F*_ST_) from the Popoolation2 software suite (Kofler et al., 2011). A minimum and maximum coverage were determined based off the mode depth +/− half the mode as per Kurland et al (2019,) and a minimum overall count of the minor allele of 3 for each pool was specified (Figure S1). Note *F*_ST_ windows were correlated to FET P-values (Figure S2). We corrected for multiple testing by using an FDR correction (see Benjamini & Hochberg 1995) and set a conservative α (1.0e-7). Manhattan plots were generated using a custom R script to plot the distribution of outliers by position and scaffold throughout the entire WTD genome. Bedtools v2.27.1 was used to characterize each window being within 25 kb up/downstream of a gene (i.e. regulatory). Numerous candidate genes for body size and head gear have been identified in ruminants (Bouwman et al., 2018; Ker et al., 2018; Wang et al., 2019) and were also complied into an *a priori* list (Table S4). For these *a priori* genes and outlier windows we identified highly differentiated SNPs using the modified chi-square test from Spitzer et al. (2020) that accounts for overdispersion.

### Transposable Elements

We used consensus transposable element (TE) sequences from the repbase database for cow to repeat mask the WTD reference genome, and subsequently merged this masked genome with the repbase TE reference sequences (Bao et al., 2015). We used the repbase database consensus sequences to re-mask the genome due to more robust information for TE identities, family and order required in this analysis. Trimmed reads for both antler and body size phenotypes were aligned to the TE-WTD merged reference genome using bwa-mem as stated in previous methods. We used the recommended workflow for the software suite popoolationTE2 v1.10.04 (Kofler et al., 2016) to create a list of predicted TE insertions. From this we calculated TE frequency differences between large and small phenotypes as well as proximity to genic regions. Here, we only examined TEs that were within 25-kb up or downstream from a gene and within the 95^th^ percentile for absolute frequency differences to allow for more features to be assessed subsequently.

### GO Pathways

To identify shared gene pathways among outlier regions we used an analysis of gene ontology (GO) terms. The program Gowinda v1.12 (Kofler & Schlötterer, 2012) was used to determine GO term enrichment, while also accounting for biases in gene length but requires input of individual SNPs. A gtf version of our annotation was created by removing duplicate genes and retaining only the longest gene – resulting in 15,395 unique genes. All SNPs in outlier windows that were also within 25-kb of a gene were included in this analysis, and compared to all SNPs in every qualifying window for antler and body size phenotypes independently. We used the program REVIGO (Supek et al., 2011) to remove redundant GO terms and to visualize semantic similarity-based scatterplots. Results from Gowinda with p-values <0.05 were used with REVIGO to generate plots of significant GO terms for biological processes of antler and body size phenotypes.

### Validation of outliers

We selected three SNPs for qPCR validation that had significant chi-square values and passed quality control with respect to primer design. Custom genotyping assays using rhAMP chemistry (Integrated DNA Technologies) were designed and genotyped on the QuantStudio 3 (Thermo Fisher Scientific). Oligo sequences and reaction parameters are provided in Table S3. Using the phenotypic category as the binary response variable, we ran a logistic regression treating the genotypic data as additive (e.g. 0-2 LA/LB alleles per locus, per individual).

## Results

### Phenotypes

The distribution of phenotypes from all pools of individuals is shown in Figure 2. We only selected the top 150 individuals at the tail ends of the distribution for measurements used in our antler and body size rankings which are representative of the extreme phenotypes. Artist renderings and the distribution of measurements for the number of antler points, beam diameter, and body length between the groups of individuals representing each extreme phenotype pool (LA, SA, LB, SB) are shown in Figure 2a. There were no differences in mean age between the pools (Figure 2b).

### Detecting genetic variants and their genomic regions

The total reads generated from resequencing are observed in Table S2 (BioProject ID PRJNA576136). The WTD genome that has a N50 of 17 MB, 727 scaffolds >50 kb (98.85% of genome), and BUSCO completeness of 91% across 303 orthologs. The genome annotation resulted in 20,750 genes. The mean allele frequency difference across the antler outlier windows were calculated to be 14% with 5249 windows meeting the FDR and α threshold (*p* <= 1×10^−7^); many of the top windows were adjacent to genes with known functions (Tables 1 & 2). For the comparison of body size, the mean allele frequency difference was 13% in 1350 windows meeting the FDR threshold.

From the list of 32 *a priori* candidate genes in the literature, we identified the highest differentiated windows (Table 3); two genes had window values exceeding the FDR (*p* < 10^−7^) for the antler analysis (OLIG1, HMGA2), and three from the body analysis (OLIG1, IGF1, TWIST1). For all individual SNPs within outlier windows and a-priori genes, the modified chi-square test with FDR correction identified 5,184 and 1,166 SNPs (p < 0.01) from the antler and body analysis, respectively.

### TE Insertions

We identified 19,160 TE insertions through the joint analysis of antler phenotype sequences (both large and small). Of these TEs, we identified 958 insertions that fell within the 95th percentile for the absolute difference between large and small phenotypes for further analysis (>0.20%; Figure S3). Of the 95th percentile TE insertions in the antler analysis, 92 were found to overlap with genes, with 283 within a 25-kb window up or downstream of genic regions. For body size, 6,610 TE insertions were identified with 332 insertions in the 95th percentile (>0.21%). Of these, 28 overlapped with genes, and 116 within 25-kb of a genic region.

### Gene Ontology Annotations

The results from Gowinda comparing all SNPs within outlier windows to background SNPs from all windows showed the top 10 enriched GO terms and gene counts (Figure S4). Six statistically enriched GO terms (FDR < 0.05) from the antler analysis were: GO:0004888 (transmembrane signaling receptor activity), GO:0004872 (receptor activity), GO:0038023 (signaling receptor activity), GO:0004930 (G-protein coupled receptor activity), GO:0022835 (transmitter-gated channel activity), GO:0022824 (transmitter-gated ion channel activity). There were no significant GO terms identified through the body analysis following FDR corrections. We show term reductions for significantly enriched GO terms (p-value <0.05) through the program REVIGO, with clustered items relating to the semantic similarity in the antler pools (Figure 4 and Figure S7).

### Validation of Outliers

We genotyped putative antler loci (RIMS1, and SRP54) and one body loci (LIRF1) split between pools which resulted in 80 and 70 individuals having complete genotype and phenotypic data, respectively (Dryad Accession no. XXXXXX). The logistic regression showed an effect of the number of antler alleles at the SRP54 locus on antler category (ß = −0.99, *p* = 0.02) and the number of LIRF1 alleles on body category (ß = −0.94, *p* = 0.03); the RIMS1 locus frequency differences wereas not validated (*p* > 0.05).

## Discussion

The study of sexually selected quantitative traits in the Anticosti Island WTD population has provided novel insights into the underlying genomic architecture of phenotypic variation. The extensive database with phenotypic measurements for these deer (n=4,466) allowed us to select individuals from the wide range of the distribution (Figure 2 & Figure S1) that are representative of extreme phenotypes for the two antler and one body size measurements. This sampling methodology is important to maximize the additive genetic variance for each trait, increasing the power and the ability to detect QTL (Kardos et al., 2016). Anticosti Island WTD form a putatively panmictic population, where we would expect gene flow across the island (Fuller et al., 2020). Consistent with this, genome-wide window *F*_ST_ was ~0.03 for both pools; however, we identified outlier regions for both traits that are atypically differentiated for what would be expected in this population (Fuller et al. 2020). The most divergent windows and TEs for antler and body traits are widely dispersed throughout the WTD genome (Figure 3), which support a polygenic model for these quantitative traits. This is consistent with the literature on body size and ornaments in mammals (Visscher et al., 2007; Bouwman et al., 2018; Wang et al., 2019).

### Genomic architecture of antlers and body size

For both traits, there is evidence for divergent windows throughout the genome that overlap with genic regions (Figure 3; Tables 1-3). Traditionally, variation in SNPs is focussed on non-synonymous variants, but it is becoming evident that non-coding regions impact phenotypic variation (Watanabe et al., 2019), often with clear relationships to promoters and enhancer regions (Pagani & Baralle 2004; Zhang & Lupski 2015; Foote et al., 2016). While some outliers are surely false positives (Whitlock & Lottheros 2015), cattle GWAS studies typically identify 100s to 1000s of QTL (Cole et al., 2011; Jiang et al., 2019). Our data suggest that it is the cumulative effects from these variants, including TE insertions, and likely pleiotropic and epistatic interactions that are ultimately driving the phenotypic variation in antler and body size in white-tailed deer.

Antlers are the only completely regenerable organ found in mammals (Li et al., 2014), a unique process that involves simultaneous exploitation of oncogenic pathways and tumor suppressor genes, and the rapid recruitment of synaptic and blood vesicles (Wang et al., 2019). Size of antlers might also be an honest signal of male quality (Vanpé et.al, 2007). Accordingly, three of the most diverged windows (Figure 3) were in close proximity to genes linked to biological processes related to development and immune responses (Table 1). LGALS9 for example is a gene that produces Galactin-9, a well studied protein expressed on tumour cells (Heusschen et.al., 2013). The function of ruminant gammadelta T cells is defined by WC1.1 (Rogers et.al., 2005), and suggests a role in host response to cell proliferation. Interestingly, MTMR2 (Myotubularin Related Protein 2) has been indicated as a candidate gene for litter size in pelibuey sheep (Hernández-Montiel et.al., 2020) and plays a role in spermatogenesis (Mruk & Cheng, 2011). Likewise, the qPCR validated SRP54 gene has been linked to bone marrow failure syndromes and skeletal abnormalities (Carapito et al., 2017). Thus, there are clear links to bone formation, oncogenic pathways, and sexual selection in the antler outlier regions.

The comparison of extreme body size phenotype sequences also revealed an array of genes of interest; however, it was more difficult than antlers to not engage in storytelling (Pavlidis et al., 2012) as traits of this nature, for example human height, involve thousands of SNPs and a multitude of biological pathways (Marouli et al., 2017). Despite the high heritability of body size in this population (Jamieson et al., 2020), there were fewer windows meeting our threshold, again consistent with many genes of small effect (see Barghi et al., 2020). Similarly, the list of 32 *a priori* genes generally revealed lower levels of variation (i.e. 29 of 32 genes had windows with *p* > 10^−7^) between phenotypes for these genes for both traits, despite their known effect; this is likely due to their influence shaping between species, rather than within species trait variation. That being said, observing three outlier genes in this non-exhaustive list does suggest focussing on candidate genes still has value in the genomic era.

### Transposable elements considerations

Traditionally, masking transposable elements reduces misalignments due to the repetitive nature of their sequences and current limitations with mapping software (Treangen & Salzberg 2012), but also misses a wealth of information that encompasses a large portion of the genome. Our focus was the frequency differences of TE insertions to properly mapped reference sequences, and how these varied between phenotypes. This stems from increasing evidence showing that TE insertions impact gene expression, and thus the variation for a given phenotype (Bourque et al., 2018). We found 283 highly divergent TE insertions in genic regions from our antler analysis and 116 from the body size analysis. The insertion of these highly variable sequences has the potential to impact gene function, and studies are now starting to emerge that both validate insertions (Lerat et al., 2018) and show evidence for positive selection (Kofler et al., 2011). This suggests the potential for TEs to modulate phenotypes within a population and we feel this is an important and often overlooked pattern in similar studies. Validating the TEs and characterizing their influence on neighbouring genes is clearly warranted given the high frequency differences between pools.

### Implications for understanding of artificial and sexual selection

Antler and body size are traits that are sexually selected for and linked to reproductive success in WTD (Newbolt et al., 2016; Morina et al., 2018), and are traits desired by hunters, managers, and farmers (e.g. antlers for velvet or large body for meat). We have highly diverged genomic regions, and thousands genome-wide variants with allele frequency differences between the phenotypic extremes for these traits. We suggest the combined effects of these variants as being the genetic drivers behind observed phenotypic variation, consistent with a polygenic model of quantitative traits (Barghi et al., 2020). Additional lines of evidence, specifically RNA-seq studies, should look at the regions identified here, but also consider interactions across the entire genomic, epigenomic, and transcriptomic landscape.

We identified and validated two of three potential QTL relating to the variation in phenotypes for WTD. While we applied the added stringency of accounting for overdispersion in the chi-square test, our inability to validate one outlier suggests a small effect size and thus requiring increased sample size (see Kardos et al. 2016), or a true type I error. Alternatively, epistatic interactions might be at play (Knief et al. 2019), but detection requires whole-genome sequencing of many individuals thereby defeating the purpose of pooled sequencing. The high effective population size and homogeneity of Anticositi Island white-tailed deer implies the need for small windows, and ultimately a SNP-based analysis. Here, expanded qPCR sample sizes might still reveal the predicted effect, and it is encouraging that approaches for low coverage pooled data are emerging (Tilk et al. 2019). As we continue to identify and validate candidate QTL, it is conceivable that a gene panel could be developed that could be used in breeding and management programs (e.g. Quality Deer Management) or assess the effects of artificial selection by trophy hunting; however until a reasonable amount of phenotypic variation can be attributed to specific QTL, a gene targeted approach for management and breeding of white-tailed deer will remain a challenge.

## Acknowledgments

This work was supported by Natural Sciences and Engineering Research Council of Canada Discovery Grant (ABAS and SDC); ComputeCanada Resources for Research Groups (ABAS); Canadian Foundation for Innovation: John R. Evans Leaders Fund (ABAS); The Symons Trust Fund for Canadian Studies (ABAS); Trent University start-up funds [ABAS]; and Industrial Chairs and Collaborative Research and Development Grants from the Natural Sciences and Engineering Research Council of Canada (SDC). We thank the outfitters of Anticosti Island and the Ministère des Forêts, de la Faune et des Parcs du Québec for logistical help associated with fieldwork, and all of the field assistants and technicians who collected samples and made phenotypic measurements throughout the duration of this project. We thank four anonymous reviewers for their extensive comments and analytical suggestions on this manuscript.

## Data Accessibility

Raw sequence data FastQ files are available on the Sequence Read Archive (Accession: PRJNA576136). All bioinformatic and analytical code available on GitLab (https://gitlab.com/WiDGeT_TrentU/PoolSeq). Raw data table have been uploaded to Dryad (Dryad Accession no. XXXXXX)

## Author Contributions

S.J.A., S.D.C., and A.B.A.S. designed the study; S.D.C., J.H.R., and S.J.A. coordinated data collection and sample curation; S.J.A. performed research and analyzed data; S.J.A. wrote the manuscript with input from J.H.R., S.D.C., and A.B.A.S.

---

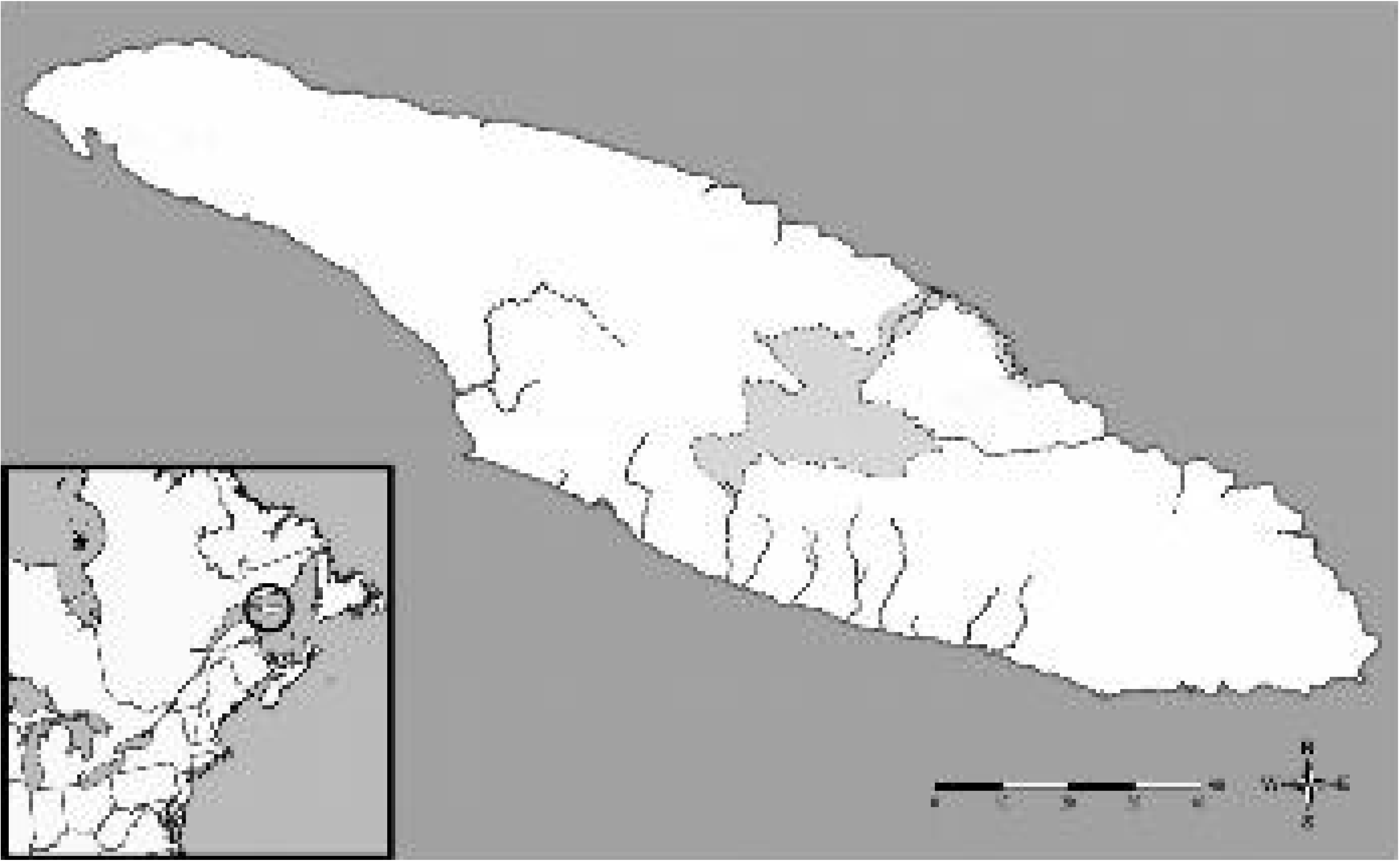

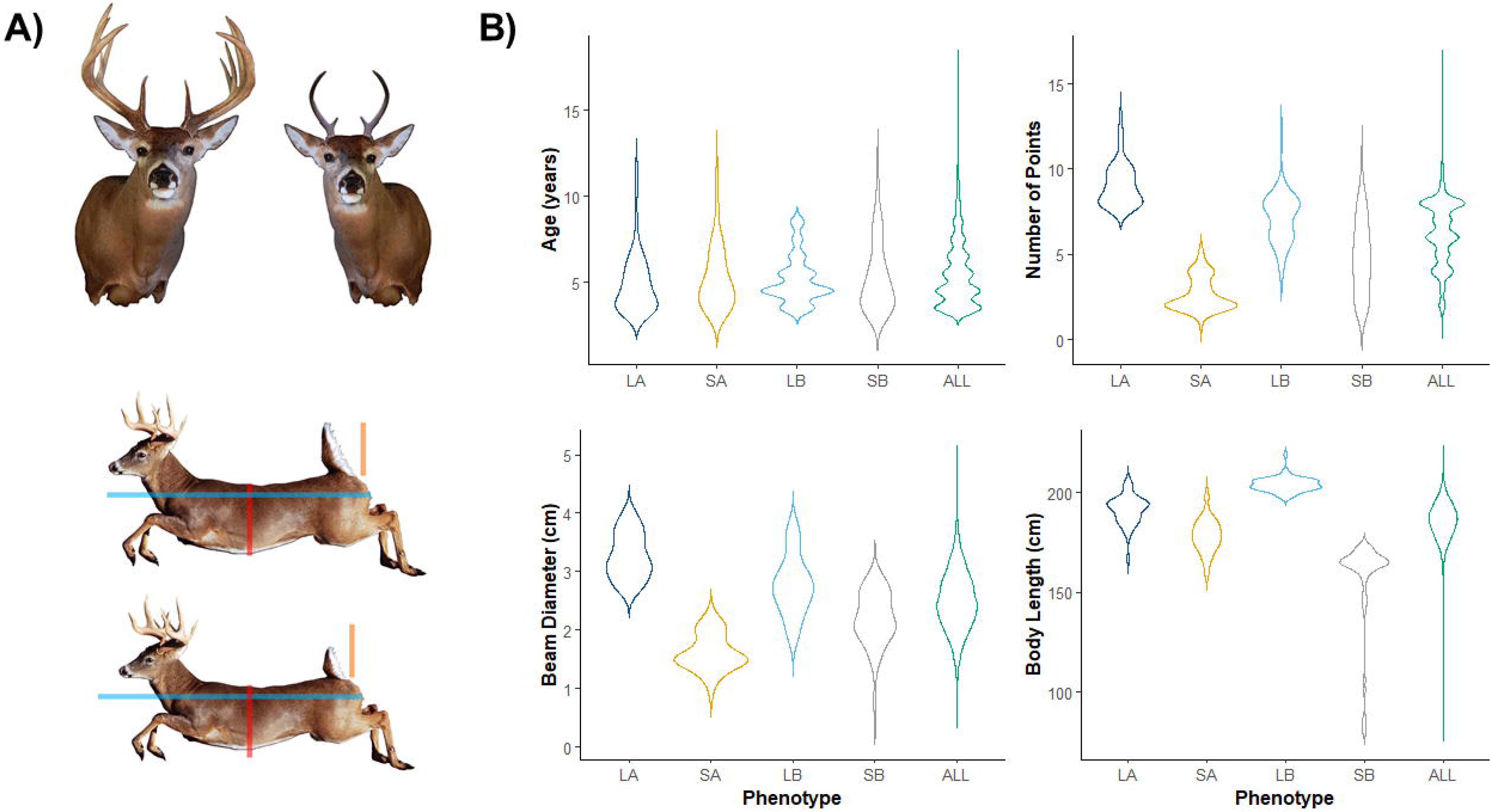

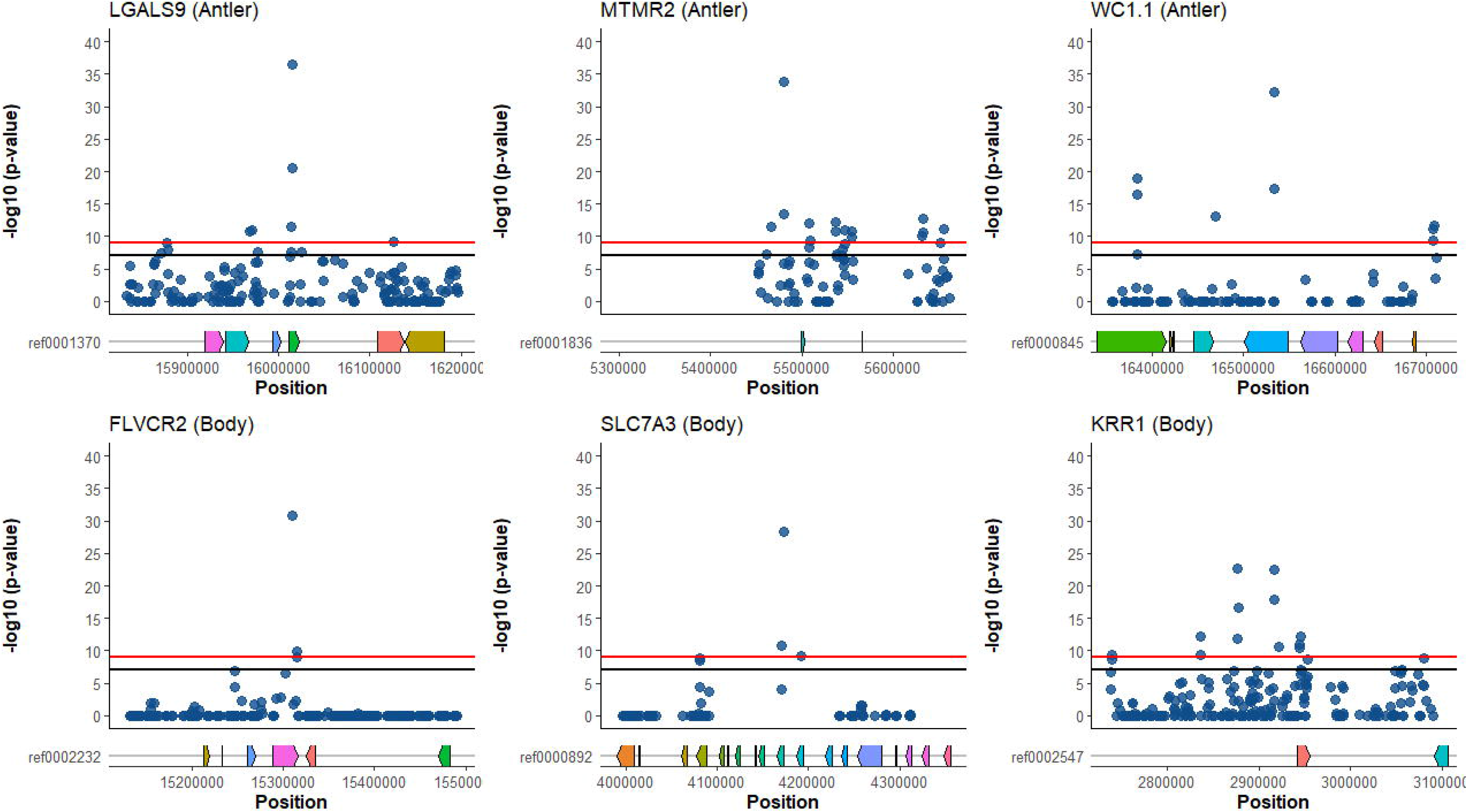

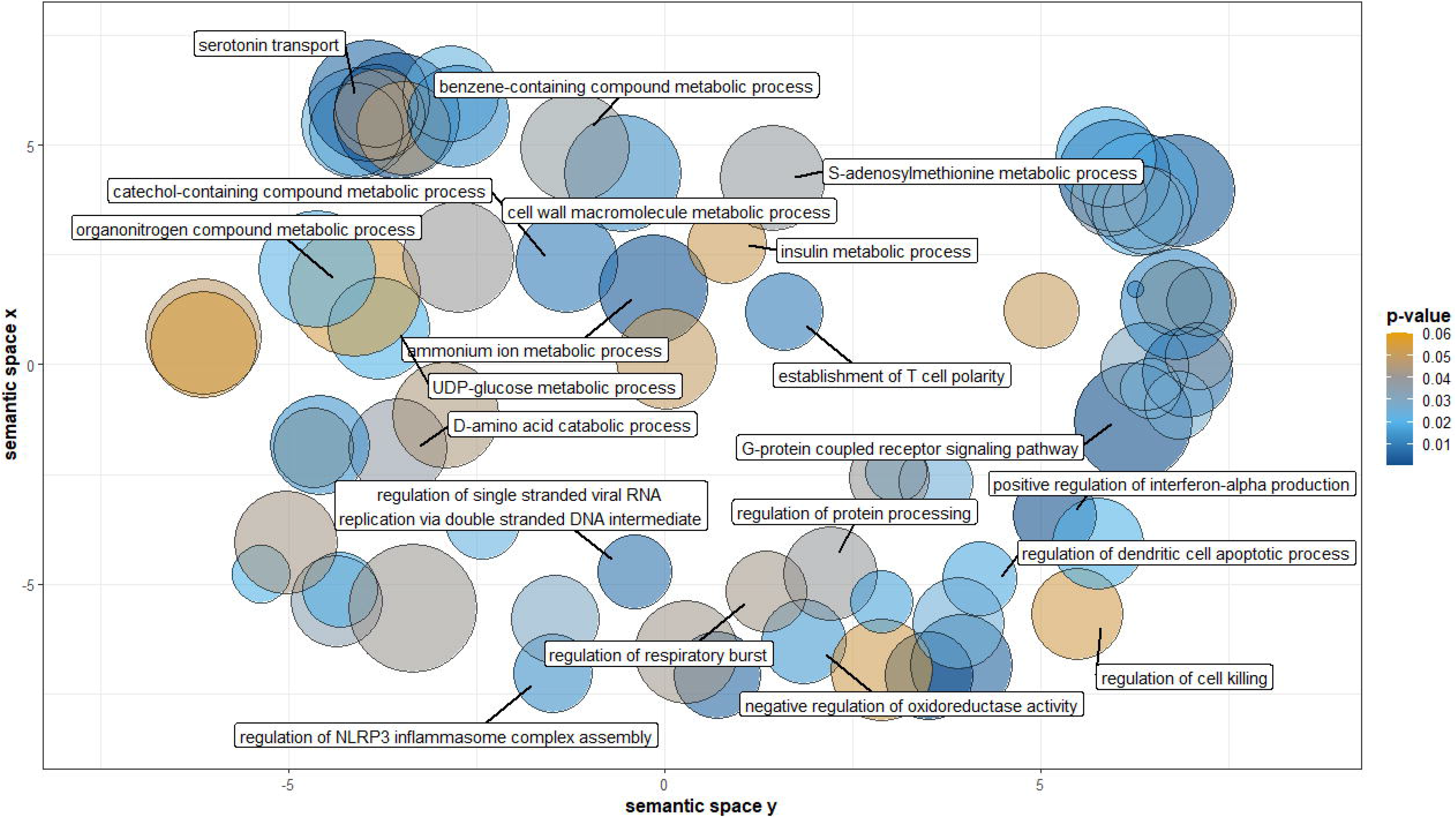

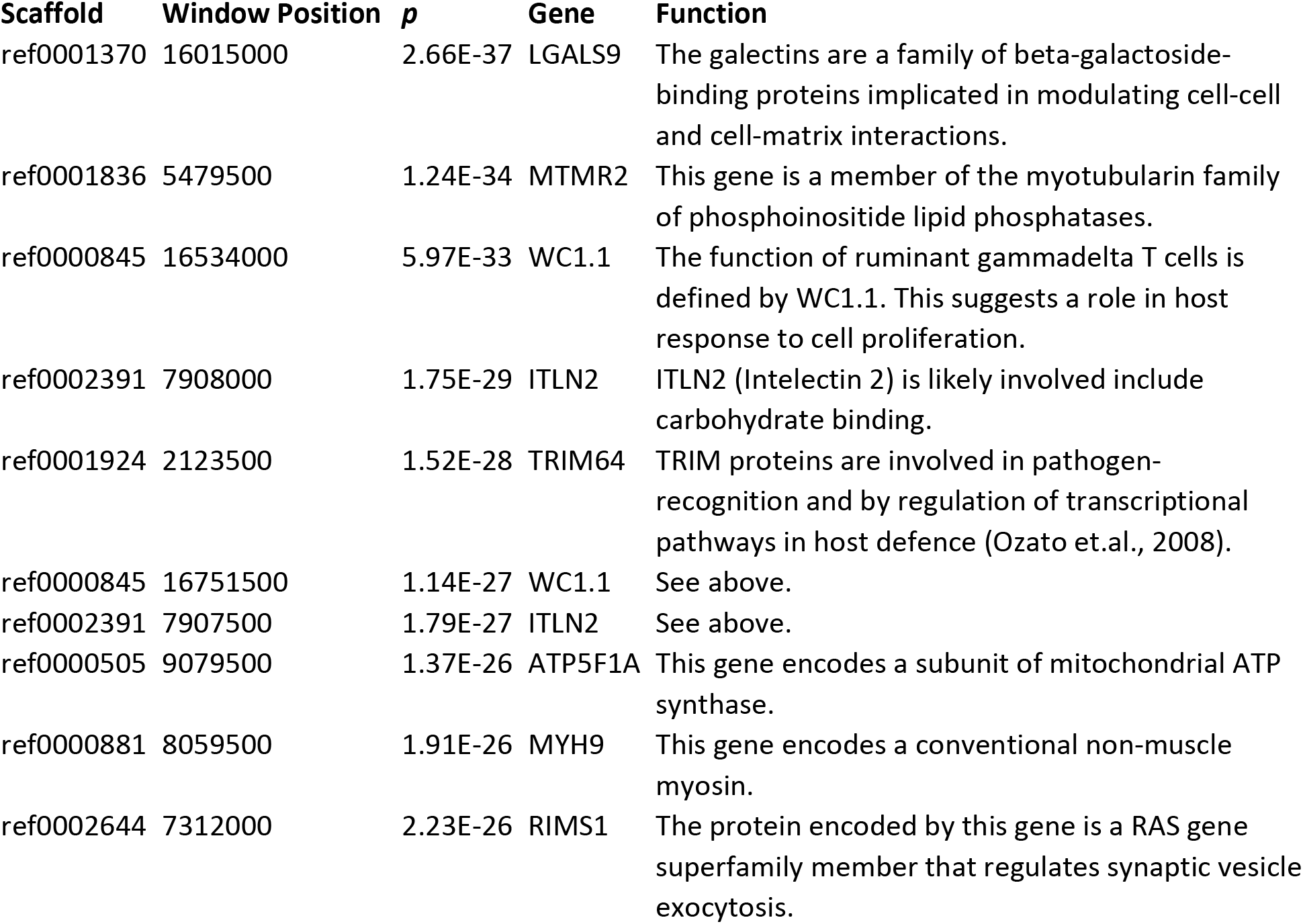
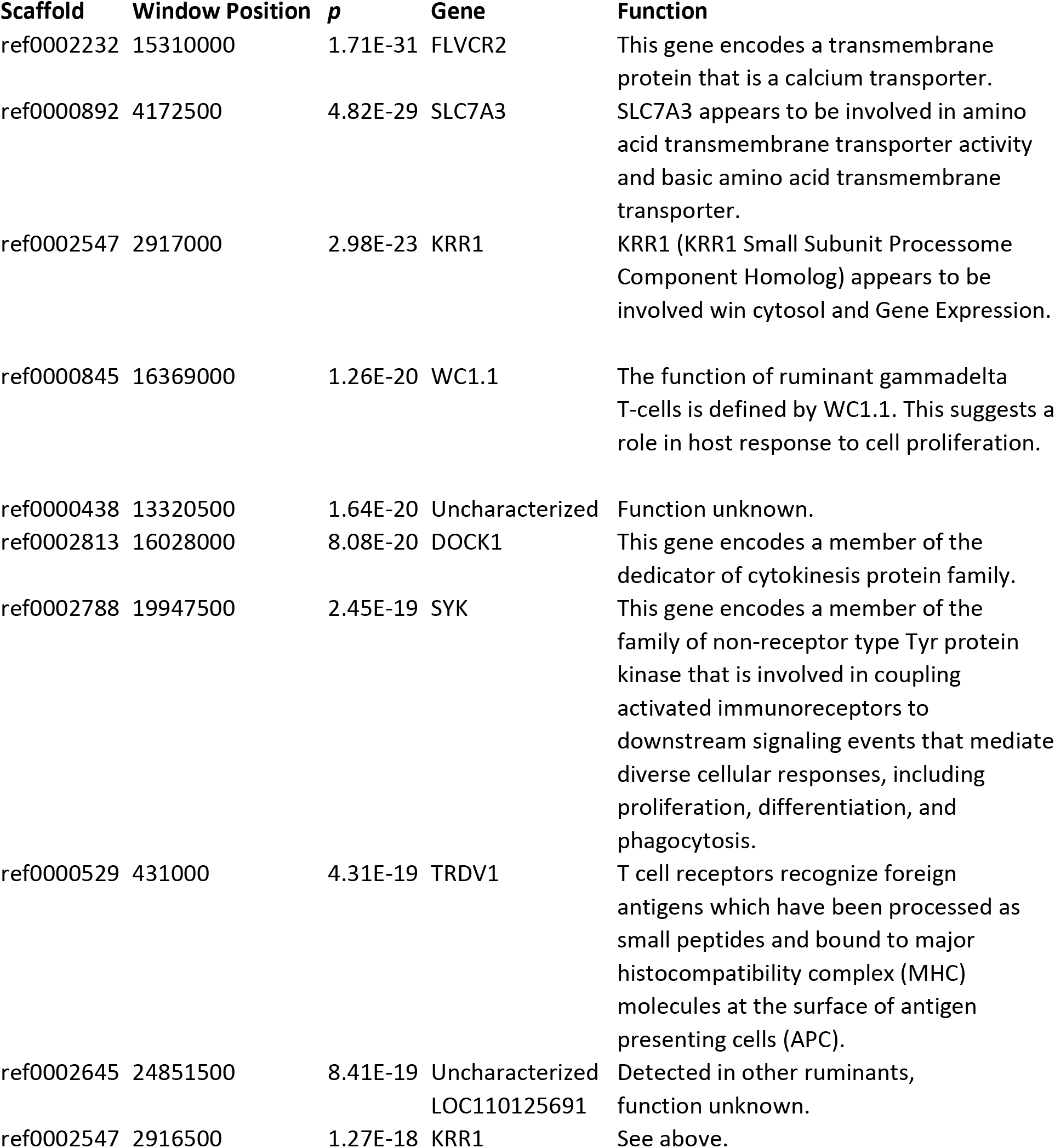

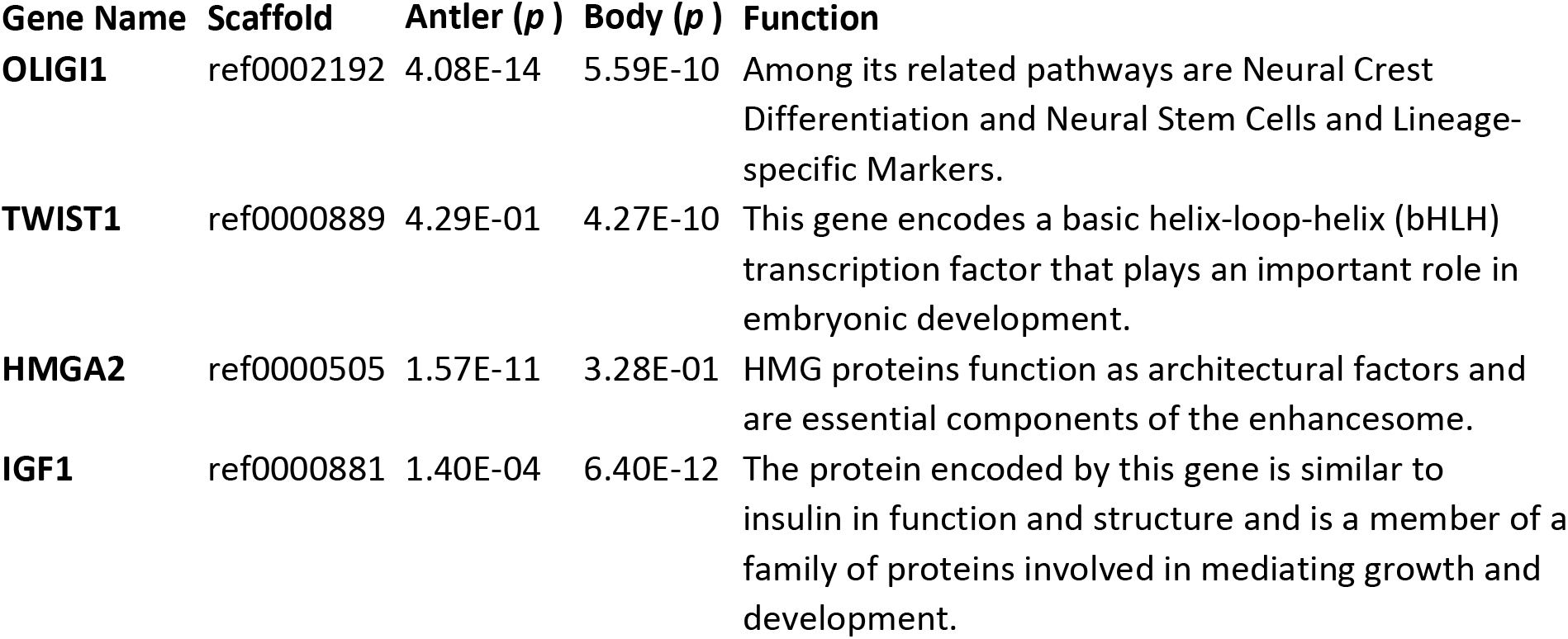

## Notes

### Competing Interest Statement

The authors have declared no competing interest.

### Summary of Updates

This version of the manuscript has been updated to reflect multiple rounds of revisions that have altered the analysis. Mainly, more rigid quality control steps have been implemented and a sliding window based analysis compared to individual SNP based. This has thus changed the representation of figures 3 & 4. The nature of the findings, however, remain consistent.

