## Supplemental Figures & Tables for "Genomic architecture of artificially and sexually selected traits in a wild cervid"

**Table 1.** The top 10 window positions from the antler trait analysis based on corrected p-values that overlapped genes or regulatory regions up/downstream. The closest gene is listed for each window. Functions of each gene are abbreviated from RefSeq/Uniprot.

| Scaffold | Window Position | *p* | Gene | Function |
| --- | --- | --- | --- | --- |
| ref0001370 | 16015000 | 2.66E-37 | LGALS9 | The galectins are a family of beta-galactoside-binding proteins implicated in modulating cell-cell and cell-matrix interactions. |
| ref0001836 | 5479500 | 1.24E-34 | MTMR2 | This gene is a member of the myotubularin family of phosphoinositide lipid phosphatases. |
| ref0000845 | 16534000 | 5.97E-33 | WC1.1 | The function of ruminant gammadelta T cells is defined by WC1.1. This suggests a role in host response to cell proliferation |
| ref0002391 | 7908000 | 1.75E-29 | ITLN2 | ITLN2 (Intelectin 2) is likely involved include carbohydrate binding. |
| ref0001924 | 2123500 | 1.52E-28 | TRIM64 | TRIM proteins are involved in pathogen-recognition and by regulation of transcriptional pathways in host defence (Ozato et.al., 2008). |
| ref0000845 | 16751500 | 1.14E-27 | WC1.1 | See above. |
| ref0002391 | 7907500 | 1.79E-27 | ITLN2 | See above. |
| ref0000505 | 9079500 | 1.37E-26 | ATP5F1A | This gene encodes a subunit of mitochondrial ATP synthase. |
| ref0000881 | 8059500 | 1.91E-26 | MYH9 | This gene encodes a conventional non-muscle myosin. |
| ref0002644 | 7312000 | 2.23E-26 | RIMS1 | The protein encoded by this gene is a RAS gene superfamily member that regulates synaptic vesicle exocytosis. |

**Table 2.** The top 10 window positions from the body trait analysis based on corrected p-values that overlapped genes or regulatory regions up/downstream. The closest gene is listed for each window. Functions of each gene are abbreviated from RefSeq/Uniprot.

| Scaffold \| Window Position \| *p* \| Gene \| Function | | | | |
| --- | --- | --- | --- | --- |
| ref0002232 | 15310000 | 1.71E-31 | FLVCR2 | This gene encodes a transmembrane protein that is a calcium transporter. |
| ref0000892 | 4172500 | 4.82E-29 | SLC7A3 | SLC7A3 appears to be involved in amino acid transmembrane transporter activity and basic amino acid transmembrane transporter |
| ref0002547 | 2917000 | 2.98E-23 | KRR1 | KRR1 (KRR1 Small Subunit Processome Component Homolog) appears to be involved win cytosol and Gene Expression. |
| ref0000845 | 16369000 | 1.26E-20 | WC1.1 | The function of ruminant gammadelta T cells is defined by WC1.1. This suggests a role in host response to cell proliferation. |
| ref0000438 | 13320500 | 1.64E-20 | Uncharacterized | Function unknown |
| ref0002813 | 16028000 | 8.08E-20 | DOCK1 | This gene encodes a member of the dedicator of cytokinesis protein family. |
| ref0002788 | 19947500 | 2.45E-19 | SYK | This gene encodes a member of the family of non-receptor type Tyr protein kinase that is involved in coupling activated immunoreceptors to downstream signaling events that mediate diverse cellular responses, including proliferation, differentiation, and phagocytosis. |
| ref0000529 | 431000 | 4.31E-19 | TRDV1 | T cell receptors recognize foreign antigens which have been processed as small peptides and bound to major histocompatibility complex (MHC) molecules at the surface of antigen presenting cells (APC). |
| ref0002645 | 24851500 | 8.41E-19 | Uncharacterized LOC110125691 | Detected in other ruminants, function unknown. |
| ref0002547 | 2916500 | 1.27E-18 | KRR1 | See above. |

**Table 3**. Genes from *A priori* list from Ker et al. (2018), Bouwman et al. (2018), and Wang et al. (2019) with identified significant associations (p < 10^-7^) to antler and body size. Represented are the scaffold, highest corrected *p-*value for windows on each gene (+/- 25-kb window) for both phenotypes and gene function abbreviated from RefSeq/Uniprot. The *a priori* list can be found in the supplemental material provided (Table S4).

| Gene Name | Scaffold | Antler (*p*) | Body (*p*) | Function |
| --- | --- | --- | --- | --- |
| OLIGI1 | ref0002192 | 4.08E-14 | 5.59E-10 | Among its related pathways are Neural Crest Differentiation and Neural Stem Cells and Lineage-specific Markers |
| TWIST1 | ref0000889 | 4.29E-01 | 4.27E-10 | This gene encodes a basic helix-loop-helix (bHLH) transcription factor that plays an important role in embryonic development. |
| HMGA2 | ref0000505 | 1.57E-11 | 3.28E-01 | HMG proteins function as architectural factors and are essential components of the enhancesome |
| IGF1 | ref0000881 | 1.40E-04 | 6.40E-12 | The protein encoded by this gene is similar to insulin in function and structure and is a member of a family of proteins involved in mediating growth and development. |

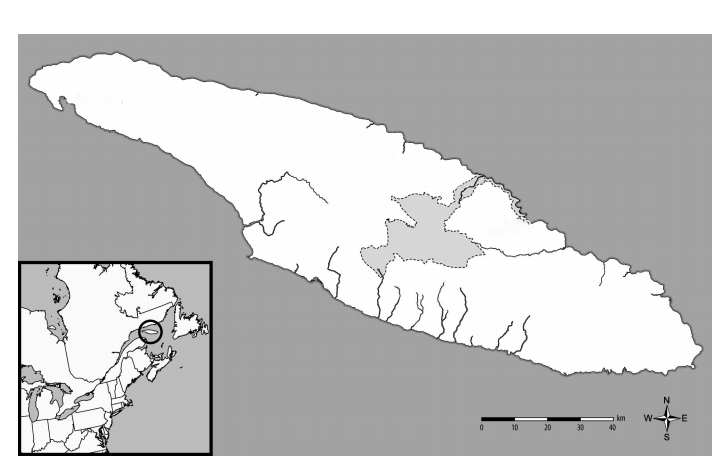

**Figure 1.** A map of Anticosti Island, Quebec where all white-tailed deer samples were acquired between 2002-2014. The grey area reflects Anticosti National Park where no hunting is permitted. The inset represents the larger North American landscape.

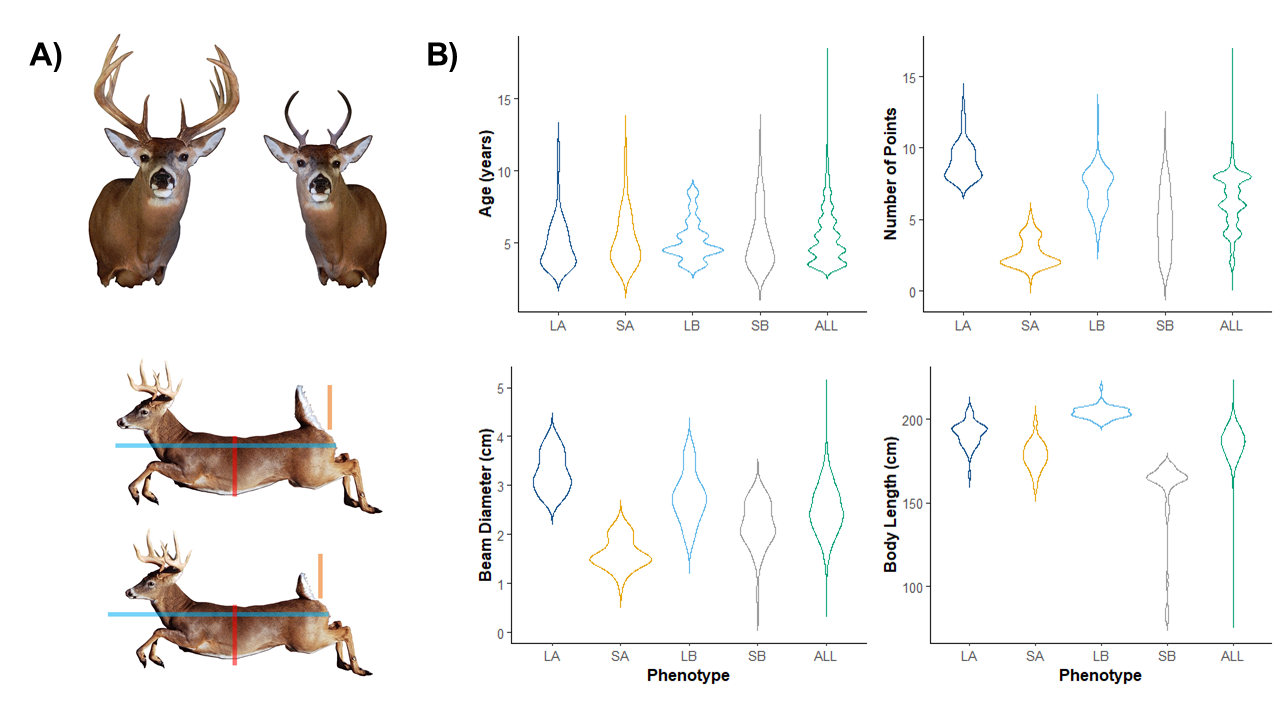

**Figure 2.** The phenotypic extremes used in our sampling methodology. a) Artist renderings of the average phenotypic measures for all individuals included in each pool. Measurements for antler pools (top) included mainbeam diameter and number of points, while body size (bottom) included body length. Scale bars are present for body size to better show differences in average body length, and chest circumference between large and small phenotypes. b) Violin plots displaying the original measurements of each individual, grouped by large antler (LA), small antler (SA), large body size (LB), and small body size (SB) pools. The phenotypes measured include age (years), number of points, beam diameter (cm), and body length (cm).

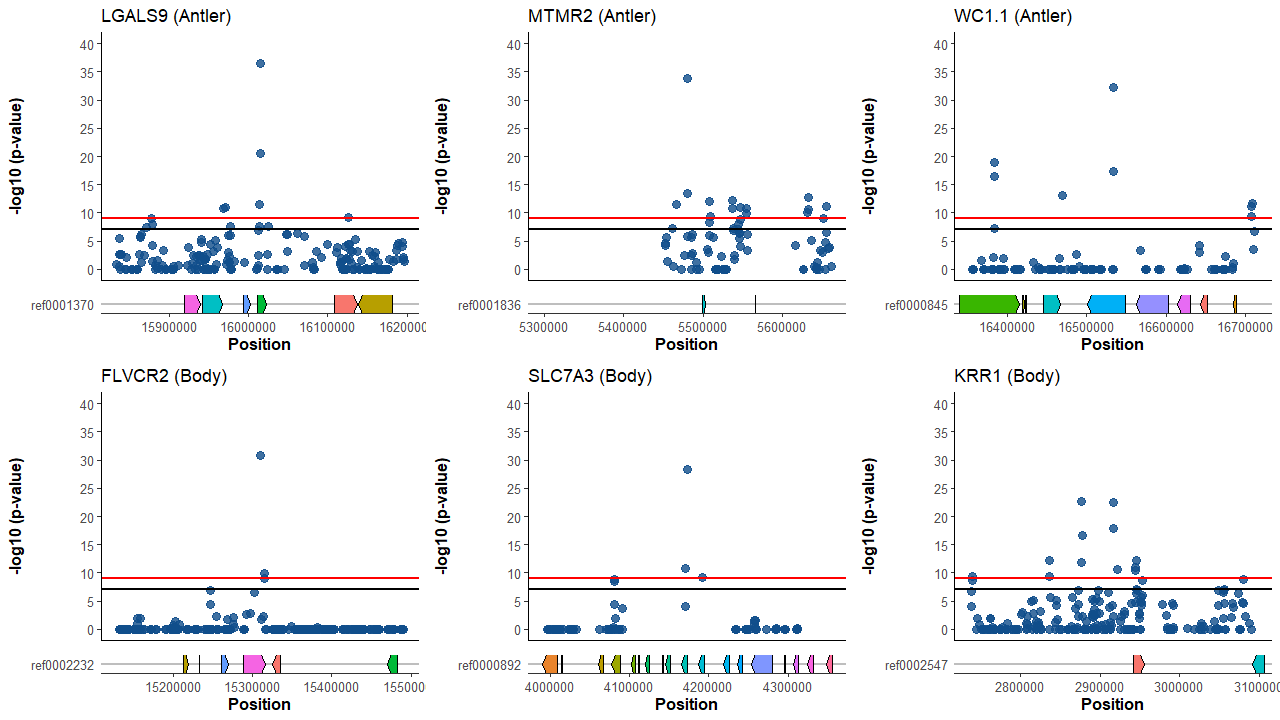

**Figure 3.** Manhattan plots representing pairwise genetic differentiation (Fisher’s Exact Test) for scaffolds of interest in 1000 bp sliding windows, with a step size of 500. Plots represent 200 kbp (+/- ) surrounding the most highly differentiated windows and their associated gene regions; the top panel representing LGALS9 (ref0001370), MTMR2 (ref0001836), and WC1.1 (ref0000845) from the comparison of antler phenotype pools and the bottom panel representing FLVCR2 (ref0002232), SLC7A3 (ref0000892), and KRR1 (ref0002547) from the comparison of body size phenotype pools. The horizontal black like represents the false detection rate (*p* = 10^-7^), while the red line represents a conservative significance threshold (*p* = 10^-9^).

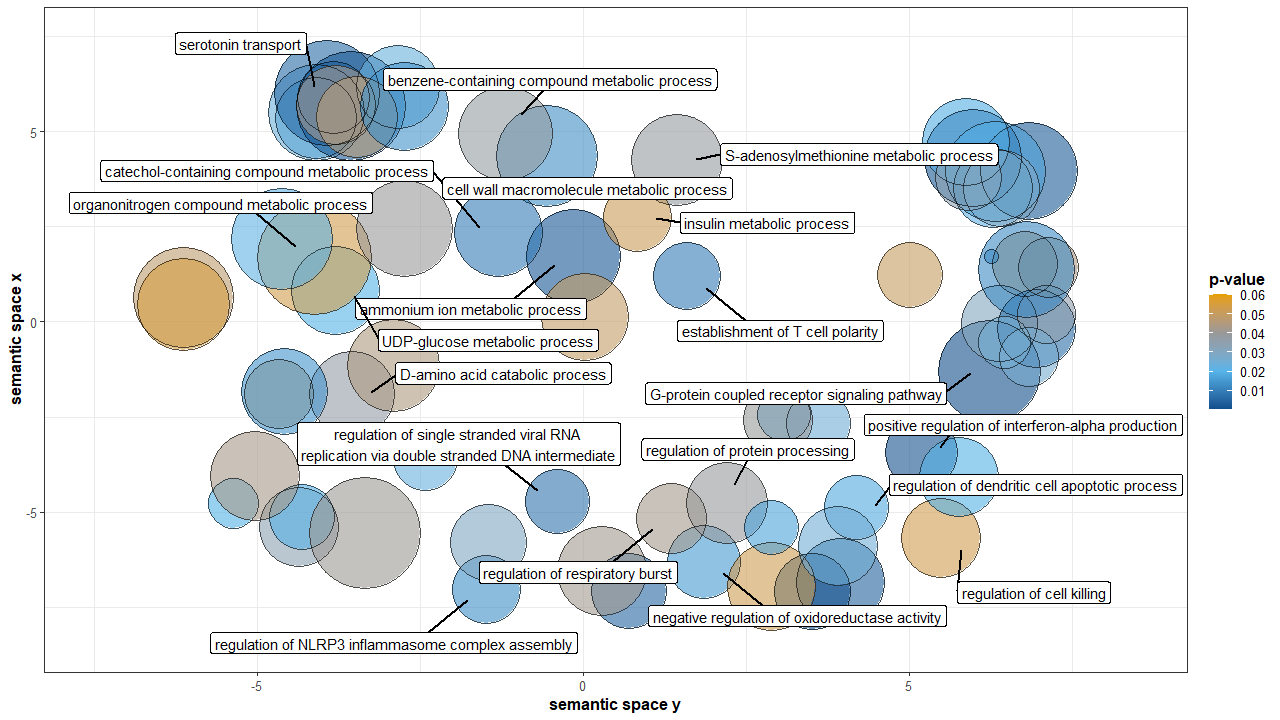

**Figure 4.** Antler analysis of GO terms grouped by semantic similarity. Points are coloured based on significance, with all terms with *p*-values < 0.05 from the output of the Gowinda analysis for gene enrichment being included in the analysis. The size of each point represents the specificity of each term; GO terms for smaller points being more specific, and larger points more general. Only points with a dispensability score > 0.30 are labeled.

**Table S1**. Mean measurements for each sequenced phenotype pool. Only measurements used in the ranking of antler and body phenotypes are present

| Phenotype Pool | Age | Body Length | Points | Beam Diameter |
| --- | --- | --- | --- | --- |
| **Large Antler** | 4.96 | 191.49 | 9.06 | 3.27 |
| **Small Antler** | 5.25 | 178.31 | 2.67 | 1.62 |
| **Large Body** | 5.17 | 204.00 | 7.13 | 2.20 |
| **Small Body** | 5.32 | 156.61 | 4.87 | 2.19 |

**Table S2**. Number of reads remaining after each filtering stepped and final genome wide coverage for each pool.

|  | Initial | Deduped | Unique | Realigned | Coverage |
| --- | --- | --- | --- | --- | --- |
| Large Antler | 733,681,602 | 683,823,491 | 586,588,595 | 586,588,595 | 33.6 |
| Small Antler | 828,746,267 | 767,420,576 | 652,843,728 | 652,843,728 | 37.4 |
| Large Body | 794,621,182 | 718,618,777 | 611,366,207 | 611,366,207 | 34.7 |
| Small Body | 824,962,249 | 728,085,328 | 618,792,299 | 618,792,299 | 35.3 |

**Table S3**. Primer sequences for the rhAMP genotyping assay used for outlier antler and body SNPs for the RIMS1, PTEN, SRP54, and LRIF1 genes.

| **Primer** | **Primer Sequence** |
| --- | --- |
| RIMS1_ASP1 | /rhAmp-F/ATCTATTATACTTAGGAAAGTTTCAAAGrGGAAA/GT4/ |
| RIMS1_ASP2 | /rhAmp-Y/ATCTATTATACTTAGGAAAGTTTCAAATrGGAAA/GT4/ |
| RIMS1_LSP | GCTCTTACCCTTTCAATCTCAGAGTGrCTAAA/GT4/ |
| SRP54_ASP1 | /rhAmp-F/ATCCTGAGCCACAGGATATrAAGGT/GT4/ |
| SRP55_ASP2 | /rhAmp-Y/ATCCTGAGCCACAGGATACrAAGGT/GT4/ |
| SRP56_LSP | GCGAGGAAATCAAGTGCTTTTACCAArGAAGA/GT3/ |
| LRIF1_ASP1 | /rhAmp-F/GCCCTTCCCTACTTAAAATGTTTrCATCA/GT2/ |
| LRIF1_ASP2 | /rhAmp-Y/GCCCTTCCCTACTTAAAATGTTCrCATCA/GT2/ |
| LRIF1_LSP | GCTAATCTGCCTTCTCTTTGGTCTrCTCAG/GT4/ |

**Table S4.** A priori list of genes from Ker et al. (2018), Bouwman et al. (2018), and Wang et al. (2019) with identified associations to antler and body size. Represented are the scaffold, highest corrected p-value for windows on each gene (+/- 25-kb window) for both phenotypes.

| Gene Name | Scaffold | Antler (*p*) | Body (*p*) | Function |
| --- | --- | --- | --- | --- |
| UHRF1 | ref0002712 | 4.24E-04 | 1.00E+00 | The protein binds to specific DNA sequences, and recruits a histone deacetylase to regulate gene expression. This gene is up-regulated in various cancers. |
| S100A10 | ref0000146 | 8.73E-01 | 1.00E+00 | S100 proteins are localized in the cytoplasm and/or nucleus of a wide range of cells, and involved in the regulation of a number of cellular processes such as cell cycle progression and differentiation |
| OLIG1 | ref0002192 | 4.08E-14 | 5.59E-10 | Among its related pathways are Neural Crest Differentiation and Neural Stem Cells and Lineage-specific Markers |
| OTOP3 | ref0000440 | 1.25E-05 | 1.00E+00 | Proton-selective channel that specifically transports protons into cells. Proton-selective channel activity is probably required in cell types that use changes in intracellular pH for cell signaling or to regulate biochemical or developmental processes. |
| HOXD | ref0002253 | 1.18E-02 | 3.20E-05 | HOXD (Homeobox D Cluster) is a Gene Cluster |
| SNAI2 | ref0002419 | 1.00E-05 | 1.71E-02 | The encoded protein acts as a transcriptional repressor that binds to E-box motifs and is also likely to repress E-cadherin transcription in breast carcinoma. This protein is involved in epithelial-mesenchymal transitions and has antiapoptotic activity |
| TWIST1 | ref0000889 | 4.29E-01 | 4.27E-10 | This gene encodes a basic helix-loop-helix (bHLH) transcription factor that plays an important role in embryonic development. |
| SOX9 | ref0000440 | 2.35E-04 | 2.80E-06 | The protein encoded by this gene recognizes the sequence CCTTGAG along with other members of the HMG-box class DNA-binding proteins. It acts during chondrocyte differentiation and, with steroidogenic factor 1, regulates transcription of the anti-Muellerian hormone (AMH) gene. Deficiencies lead to the skeletal malformation syndrome campomelic dysplasia, frequently with sex reversal. |
| RXFP2 | ref0002699 | 2.79E-02 | 3.01E-02 | This gene encodes a member of the GPCR (G protein-coupled, 7-transmembrane receptor) family |
| SOX10 | ref0000845 | 3.00E-05 | 6.02E-02 | This gene encodes a member of the SOX (SRY-related HMG-box) family of transcription factors involved in the regulation of embryonic development and in the determination of the cell fate. This protein acts as a nucleocytoplasmic shuttle protein and is important for neural crest and peripheral nervous system development. |
| NGFR | ref0000440 | 1.54E-03 | 1.71E-05 | Nerve growth factor receptor contains an extracellular domain. |
| FOS | ref0002232 | 1.07E-02 | 1.00E+00 | As such, the FOS proteins have been implicated as regulators of cell proliferation, differentiation, and transformation. In some cases, expression of the FOS gene has also been associated with apoptotic cell death. |
| REL | ref0001545 | 1.00E+00 | 1.00E+00 | Members of this family regulate genes involved in apoptosis, inflammation, the immune response, and oncogenic processes. This proto-oncogene plays a role in the survival and proliferation of B lymphocytes. |
| FAM83A | ref0002419 | 6.66E-07 | 4.42E-02 | Probable proto-oncogene that functions in the epidermal growth factor receptor/EGFR signaling pathway. |
| PML | ref0002500 | 9.72E-03 | 6.14E-02 | The protein encoded by this gene is a member of the tripartite motif (TRIM) family. The TRIM motif includes three zinc-binding domains, a RING, a B-box type 1 and a B-box type 2, and a coiled-coil region. This phosphoprotein localizes to nuclear bodies where it functions as a transcription factor and tumor suppressor. Its expression is cell-cycle related and it regulates the p53 response to oncogenic signals. The gene is often involved in the translocation with the retinoic acid receptor alpha gene associated with acute promyelocytic leukemia (APL). Extensive alternative splicing of this gene results in several variations of the protein's central and C-terminal regions; all variants encode the same N-terminus. Alternatively spliced transcript variants encoding different isoforms have been identified. |
| NMT2 | ref0001234 | 5.07E-05 | 1.39E-02 | This gene encodes one of two N-myristoyltransferase proteins. N-terminal myristoylation is a lipid modification that is involved in regulating the function and localization of signaling proteins. The encoded protein catalyzes the addition of a myristoyl group to the N-terminal glycine residue of many signaling proteins, including the human immunodeficiency virus type 1 (HIV-1) proteins, Gag and Nef. |
| CD2AP | ref0001640 | 1.00E+00 | 1.00E+00 | This gene encodes a scaffolding molecule that regulates the actin cytoskeleton. |
| ELOVL6 | ref0000086 | 1.25E-03 | 7.35E-03 | Uses malonyl-CoA as a 2-carbon donor in the first and rate-limiting step of fatty acid elongation |
| S100A8 | ref0002511 | 1.24E-01 | 2.23E-01 | S100 proteins are localized in the cytoplasm and/or nucleus of a wide range of cells, and involved in the regulation of a number of cellular processes such as cell cycle progression and differentiation. |
| ISG15 | - | 1.00E+00 | 1.00E+00 | Several functions have been ascribed to the encoded protein, including chemotactic activity towards neutrophils, direction of ligated target proteins to intermediate filaments, cell-to-cell signaling, and antiviral activity during viral infections. |
| CNOT3 | ref0000892 | 1.00E+00 | 5.25E-03 | May be involved in metabolic regulation; may be involved in recruitment of the CCR4-NOT complex to deadenylation target mRNAs involved in energy metabolism. |
| CCDC69 | ref0001167 | 9.72E-05 | 6.62E-07 | May act as a scaffold to regulate the recruitment and assembly of spindle midzone components. |
| IGF2 | ref0002556 | 1.00E+00 | 1.00E+00 | This gene encodes a member of the insulin family of polypeptide growth factors, which are involved in development and growth. |
| HMGA2 | ref0000505 | 1.57E-11 | 3.28E-01 | HMG proteins function as architectural factors and are essential components of the enhancesome |
| CCND2 | ref0000845 | 3.12E-03 | 1.00E+00 | The protein encoded by this gene belongs to the highly conserved cyclin family, whose members are characterized by a dramatic periodicity in protein abundance through the cell cycle. Cyclins function as regulators of CDK kinases. Different cyclins exhibit distinct expression and degradation patterns which contribute to the temporal coordination of each mitotic event. This cyclin forms a complex with CDK4 or CDK6 and functions as a regulatory subunit of the complex, whose activity is required for cell cycle G1/S transition. This protein has been shown to interact with and be involved in the phosphorylation of tumor suppressor protein Rb. Knockout studies of the homologous gene in mouse suggest the essential roles of this gene in ovarian granulosa and germ cell proliferation. High level expression of this gene was observed in ovarian and testicular tumors. Mutations in this gene are associated with megalencephaly-polymicrogyria-polydactyly-hydrocephalus syndrome 3 |
| LCORL | ref0001631 | 1.00E+00 | 1.00E+00 | This gene encodes a transcription factor that appears to function in spermatogenesis. Polymorphisms in this gene are associated with measures of skeletal frame size and adult height. Alternative splicing results in multiple transcript variants. |
| NCAPG | ref0001631 | 1.00E+00 | 1.00E+00 | This gene encodes a subunit of the condensin complex, which is responsible for the condensation and stabilization of chromosomes during mitosis and meiosis. |
| PLAG1 | ref0002419 | 4.39E-02 | 1.00E+00 | PLAG1, which is developmentally regulated, has been shown to be consistently rearranged in pleomorphic adenomas of the salivary glands. |
| TNNI2 | ref0002556 | 2.18E-05 | 1.12E-02 | This gene encodes a fast-twitch skeletal muscle protein, a member of the troponin I gene family, and a component of the troponin complex including troponin T, troponin C and troponin I subunits. The troponin complex, along with tropomyosin, is responsible for the calcium-dependent regulation of striated muscle contraction. |
| TCP11 | ref0002244 | 1.24E-05 | 1.00E+00 | Plays a role in the process of sperm capacitation and acrosome reactions. Probable receptor for the putative fertilization-promoting peptide (FPP) at the sperm membrane that may modulate the activity of the adenylyl cyclase cAMP pathway |
| IGF1 | ref0000881 | 1.40E-04 | 6.40E-12 | The protein encoded by this gene is similar to insulin in function and structure and is a member of a family of proteins involved in mediating growth and development. |

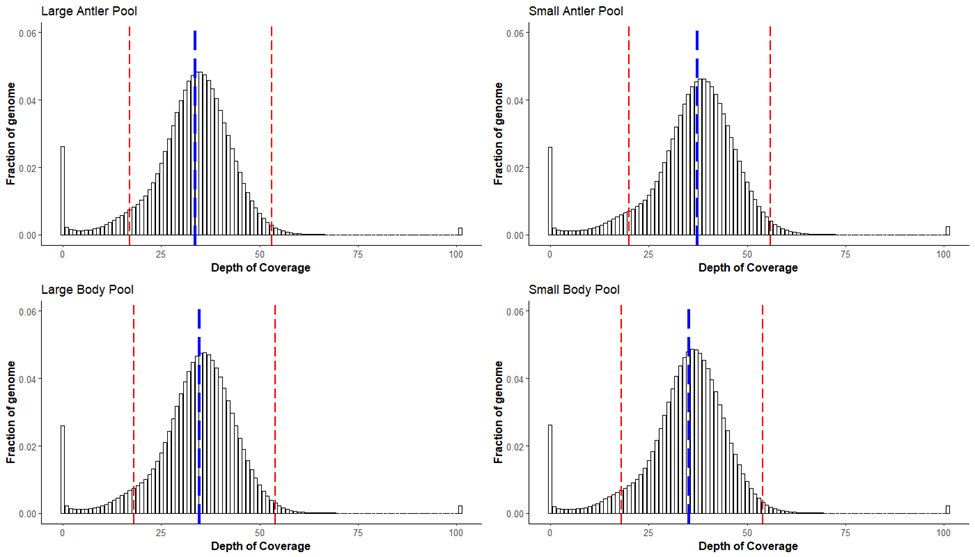

**Figure S1.** Depth of genome coverage throughout the white-tailed deer genome for all pools in our analysis. Dotted blue lines represent the calculated mean depth, while dotted red lines represent the upper and lower cut off values (+/- half the average mode from each pool).

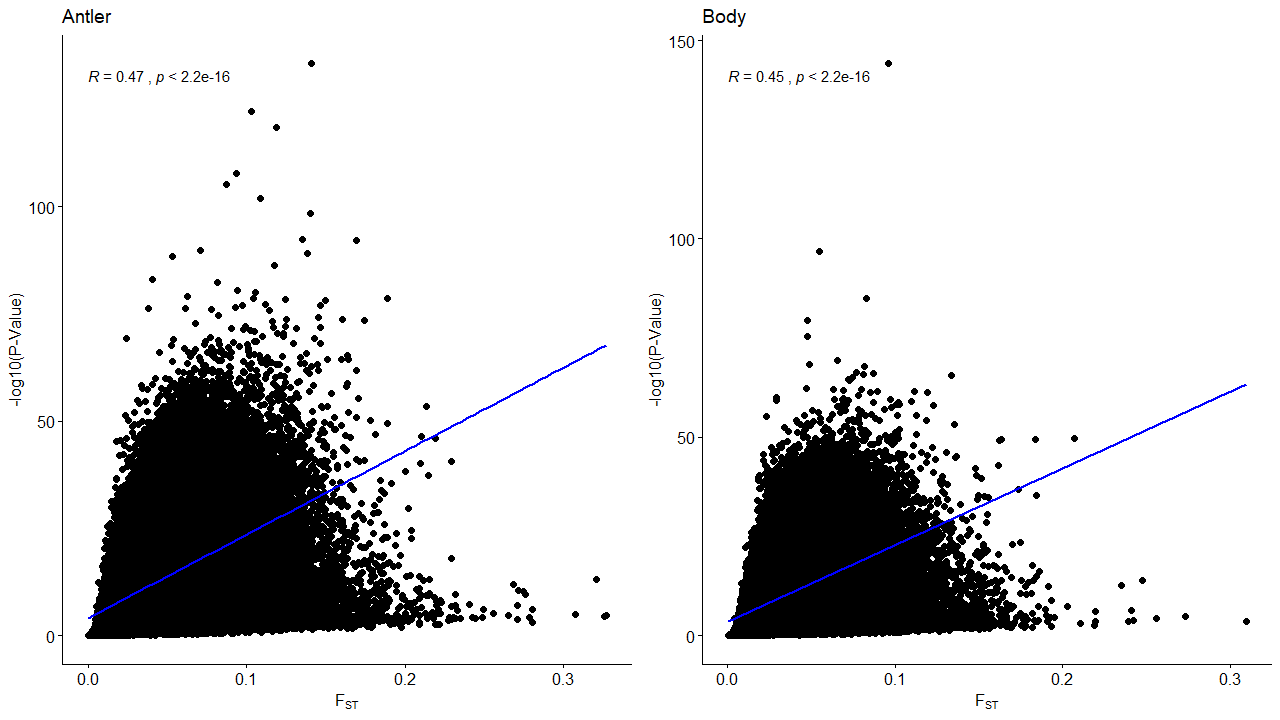

**Figure S2**. Pearson correlation between *F*_ST_ and FET values in 1000 bp sliding windows (500 bp step size) for all SNPs in both antler (left) and body (right) analysis. Analysis shows a positive correlation (Antler, Pearson’s R = 0.47, p < 2.2e-16; Body, Pearson’s R = 0.45, p < 2.2e-16).

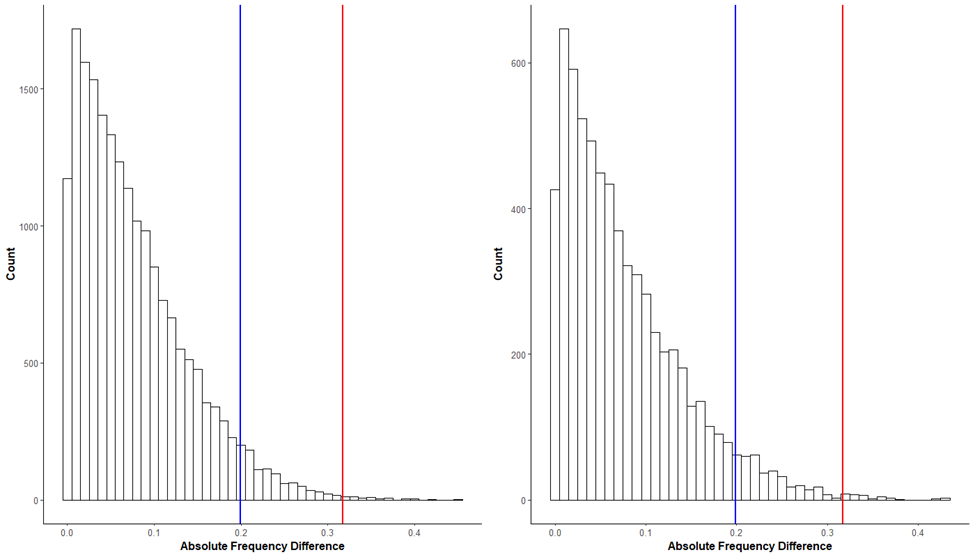

**Figure S3**. Histogram of the absolute frequency difference between transposable elements obtained through comparison of large and small antler pool analysis. Vertical blue bar represents the 95^th^ percentile, and vertical red bar represents the 99^th^ percentile.

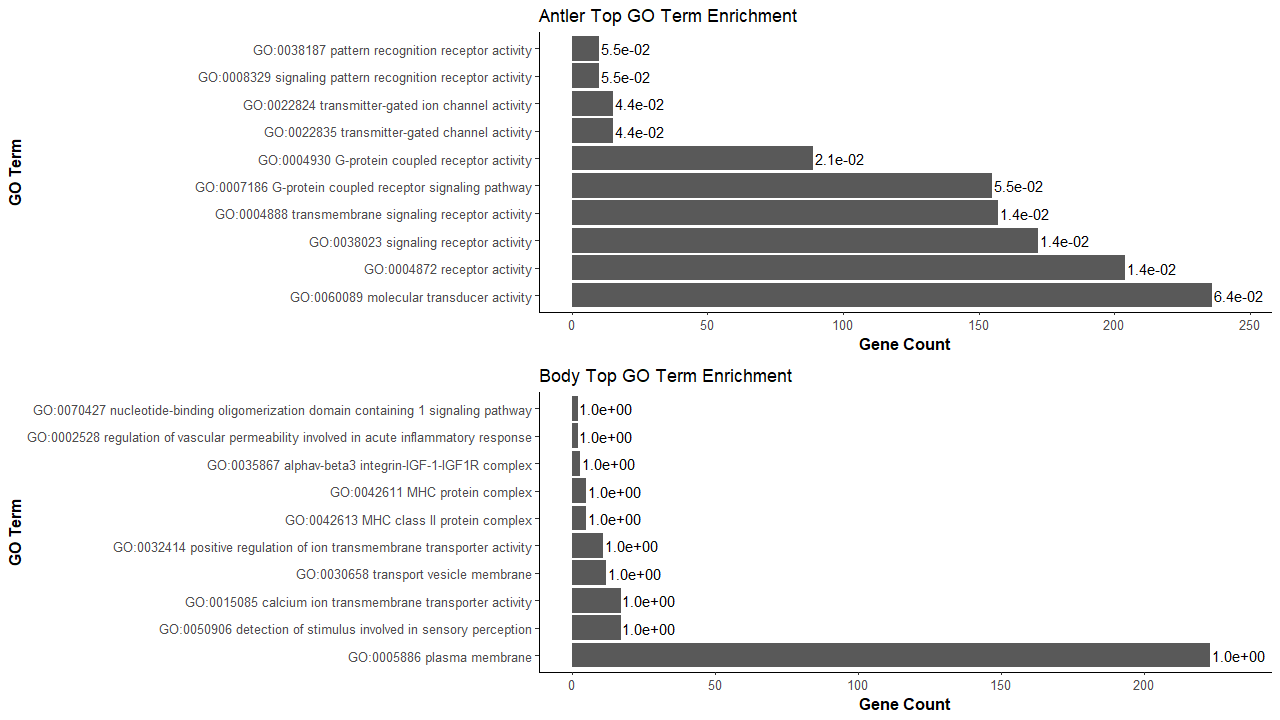

**Figure S4.** Histograms of the top ten enriched GO terms identified through outlier genes in the antler (top) and body (bottom) genome wide association study.

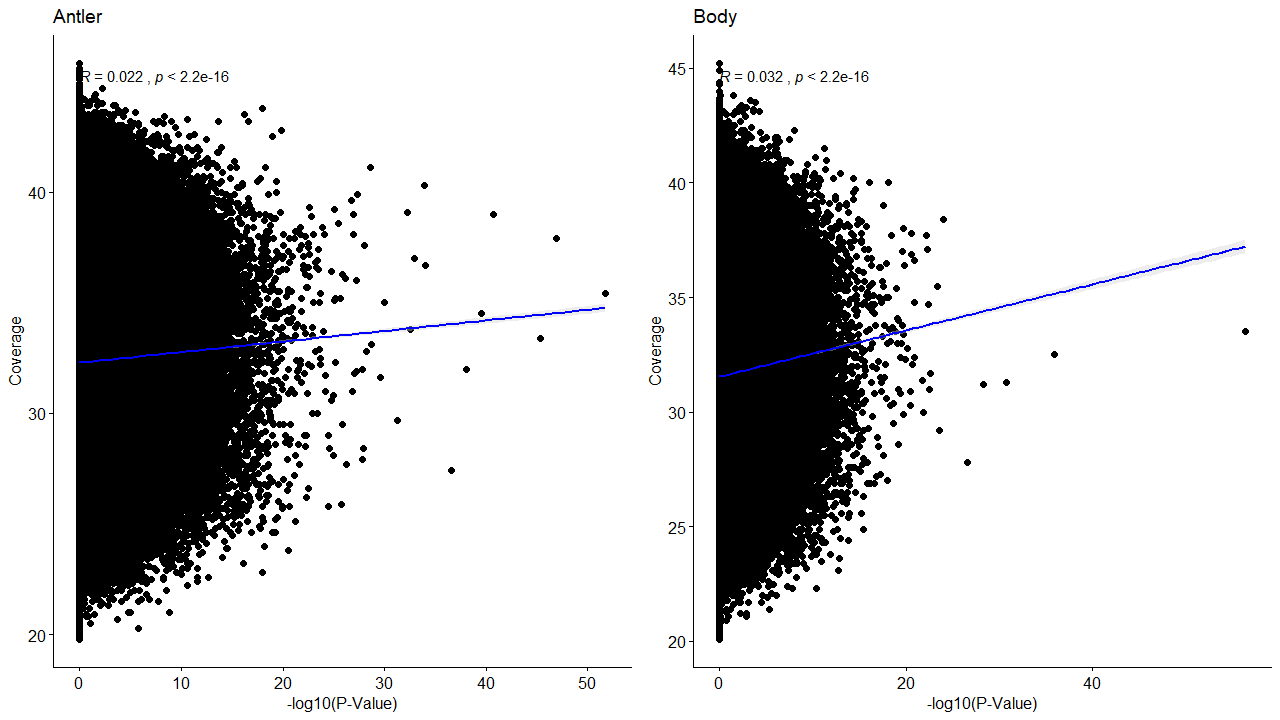

**Figure S5.** Scatter plot of window coverage vs *F*_ST_ for antler (left) and body (right) analysis. Analysis shows a slight positive correlation (Antler, Pearson’s R = 0.022, p < 2.2e^-16^; Body, Pearson’s R = 0.032, p < 2.2e^-16^).

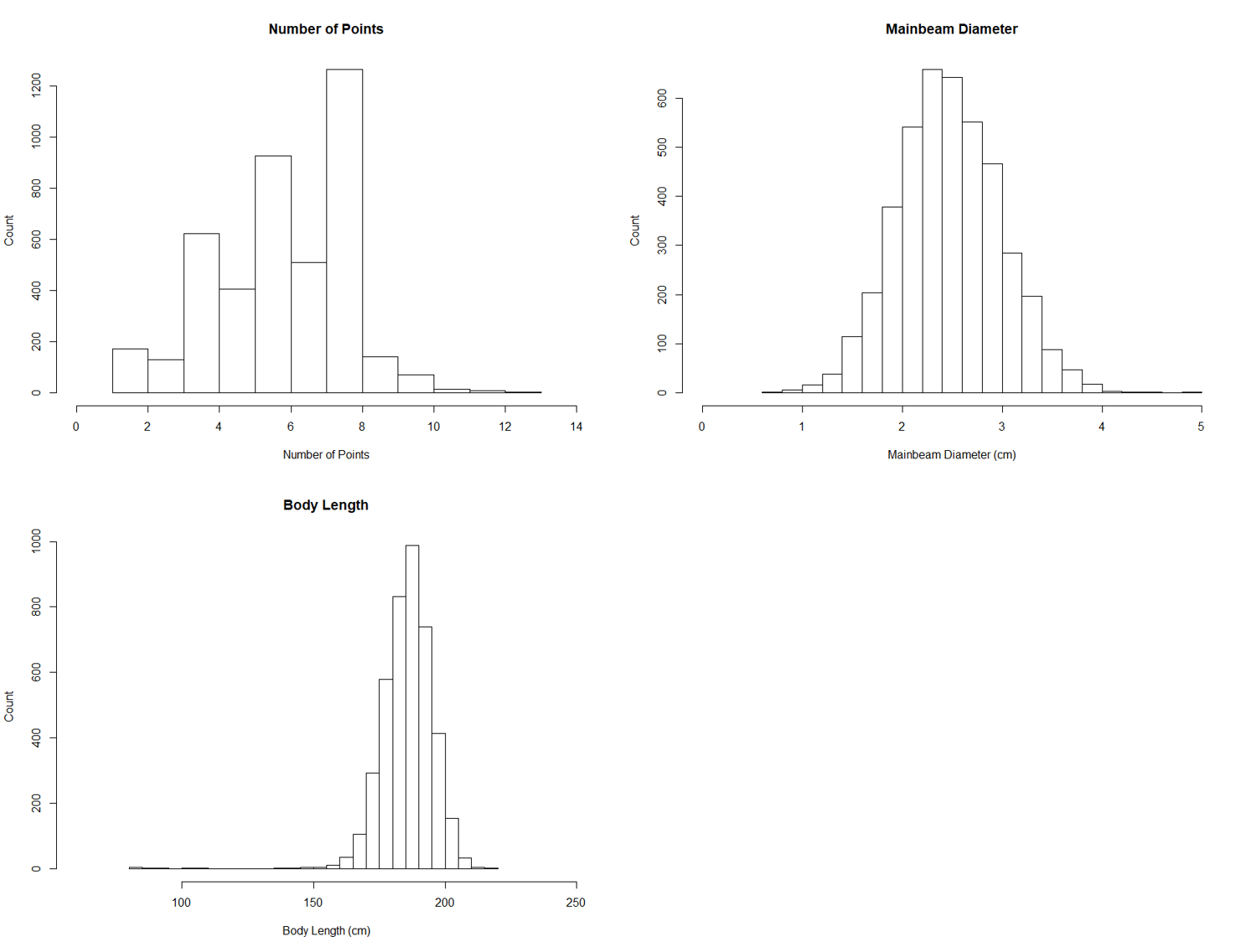

**Figure S6.** Histogram for the count of individual measurements for the number of antler points (top left), mainbeam diameter (top right) and body length (bottom left) from the entire white-tailed deer database (n=4,466).

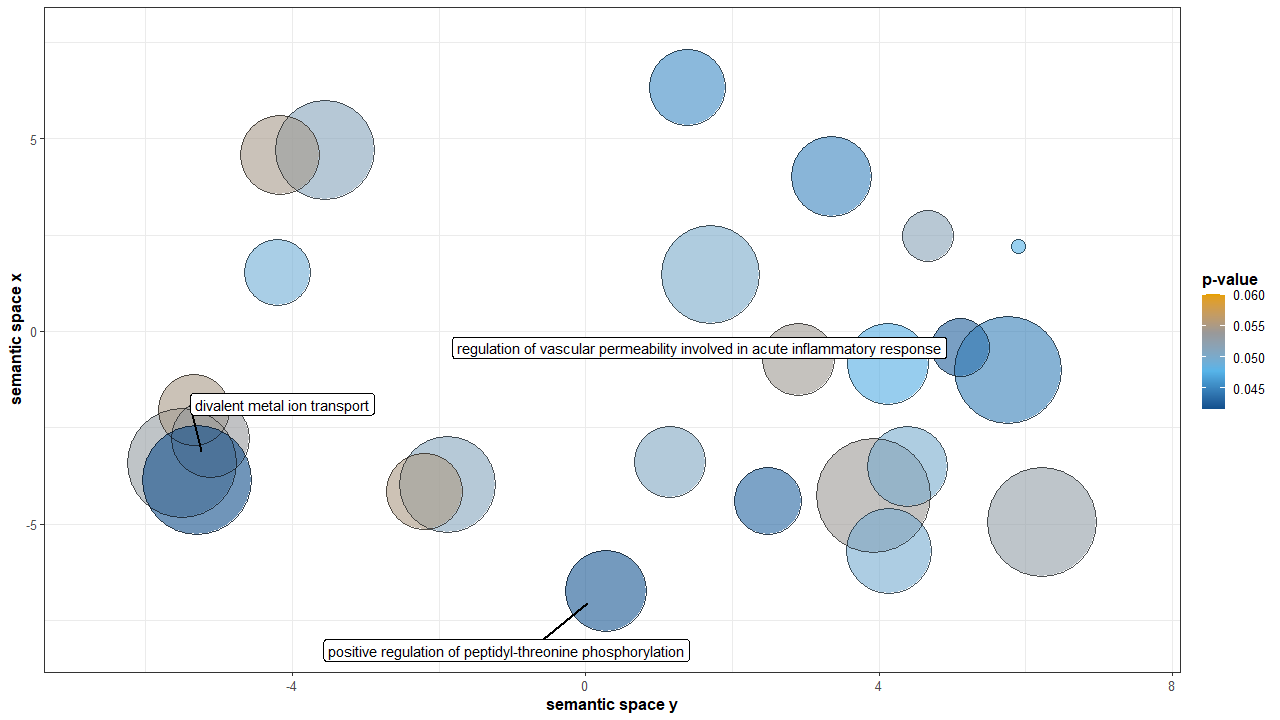

**Figure S7.** Body Analysis of GO terms grouped by semantic similarity. Points are coloured based on significance, with all terms with *p*-values < 0.05 from the output of the Gowinda analysis for gene enrichment being included in the analysis. The size of each point represents the specificity of each term; GO terms for smaller points being more specific, and larger points more general. Only points with a dispensability score < 0.30 are labeled.
